# Acute-stress impairs cytoprotective mechanisms through neural inhibition of the insulin pathway

**DOI:** 10.1101/294645

**Authors:** Maria José De Rosa, Tania Veuthey, Jeremy Florman, Jeff Grant, Gabriela Blanco, Natalia Andersen, Jamie Donnelly, Diego Rayes, Mark J. Alkema

## Abstract

Persistent activation of the “fight-or-flight” response accelerates aging and increases the susceptibility to disease. We show that repeated induction of the *C. elegans* flight response inhibits conserved cytoprotective mechanisms. This acute-stress response activates neurons that release tyramine, the invertebrate analog of adrenaline/noradrenaline. Tyramine stimulates the DAF-2/Insulin/IGF-1 pathway and precludes the nuclear translocation of the DAF-16/FOXO transcription factor through the activation of an adrenergic-like receptor TYRA-3 in the intestine. In contrast, environmental long-term stressors, such as heat or oxidative stress, reduce tyramine release allowing the induction of FOXO-dependent cytoprotective genes. These findings demonstrate how a neural stress-hormone signaling provides a state-dependent neural switch between acute and long-term stress responses, and provide mechanistic insights how acute stress impairs cellular defensive systems.

**One Sentence Summary**: The “fight-or-flight” response reduces resistance to environmental challenges.

---

Stress consists of the elicited biological response when an animal perceives a threat to its homeostasis (1, 2). The animal’s stress response requires different adaptive strategies depending on the nature and duration of the stressor. For instance, when confronted with a potentially harmful stressor the first and most efficient response is a behavioral one. An animal can avoid the negative impacts by removing itself from the threat; *e.g.* by seeking the shade if body temperatures become elevated or by fleeing to avoid predators. An attack, or even the perception of a predator elicits a rapid “fight-or-flight” response to enhance the animal’s chance of survival. In mammals, the acute fight-or-flight response leads to the activation of the sympathetic nervous system and the release of “stress hormones”, such as adrenaline and noradrenaline (3). These neurohormones trigger an increase in heart rate, blood flow, respiration and release of glucose from energy stores, which prepare the animal for vigorous muscle activity and physical exertion. The high energy-demand of a flight response is short-lived and has no significant impact on the animal’s welfare once the threat disappears. However, a behavioral response may be insufficient in some cases. Stressful situations such as temperature challenges, food shortage, hypoxia or oxidative stress generally occur on longer time scales and need more gradual and persistent strategies for adaptation (4, 5). The response to long-term environmental stressors triggers the activation of highly conserved cytoprotective processes. Stress-induced activation of transcription factors such as the Forkhead box O (FOXO), Heat Shock Factors (HSFs) and Hypoxia-Inducible Factors (HIFs), drives the expression of antioxidant enzymes and protein chaperones, to cope with protein misfolding and aggregation (6, 7). Perhaps counterintuitively, exposure to a low dose of a harmful stressor can make an organism more resistant to higher and otherwise detrimental doses of the same stressor (8). The induction of improved stress tolerance, called hormesis, has been observed in response to oxidative stress, heat and radiation, in organisms ranging from microbes to plants to humans (9, 10). This adaptive response to cellular stress, can even lead to lifespan extension and cross adaptation to other stress factors (11). Improved stress tolerance is thought to depend on protective activation of cytoprotective mechanisms that permit a more effective response to stronger environmental stressors (8). However, the concept of hormesis, and the nature of its underlying molecular mechanisms, remains contentious (12, 13).

While moderate amounts of stress may improve health and increase lifespan (14, 15), repeated activation of the acute stress response can have negative impacts. For instance repeated exposure to predators increases oxidative stress, reduces disease resistance and shortens lifespan in insects (16). In mammals, the chronic activation of the fight-or-flight response results in immunosuppression (17). When the acute stress response is perpetuated, as is the case of patients with post-traumatic stress disorder (PTSD), it can lead to accelerated aging and increased susceptibility to metabolic, cardiovascular and infectious diseases (18, 19). The acute stress response can be triggered by an actual or perceived threat, suggesting a neuronal, non-cell autonomous regulation of cellular defense mechanisms. This is supported by studies in the nematode *Caenorhabditis elegans* in which neuronal perception of harmful situations is needed for the coordination of the systemic stress response (7, 20). However, how the nervous system affects long-term stress response in animals remains largely obscure.

Like other animals, *C. elegans* faces challenges in its environment that occur either abruptly (*e.g.* predation) or more progressively (food shortage, osmotic stress, oxidation, high or low environmental temperatures). Much like in mammals, the nematode stress response to *e.g.* heat, starvation and oxidative stress features a central role for the DAF-2/ Insulin/IGF-1 signaling (IIS) pathway and the activation of DAF-16/FOXO, SKN-1/NRF and HSF-1 transcription factors (21). In response to acute challenges *C. elegans* can engage in an escape-response where it rapidly moves away from a harmful stimulus. The escape response triggers the release of tyramine (22, 23), which is structurally and functionally related to adrenaline/noradrenaline (24). Tyramine plays a crucial role in the coordination of independent motor programs, which increases the animal’s chances to escape predation (22, 23, 25, 26).

In this study we examine how the nervous system controls and coordinates behavioral and cellular defense mechanisms. We find that sub-lethal exposure to long-term stressors inhibits tyramine signaling and protects the animal from subsequent stronger stressors. In contrast, acute-stressors trigger tyramine signaling, making animals more sensitive to long-term stressors. Tyramine activation of an adrenergic-like receptor, TYRA-3, in the intestine stimulates the DAF-2/IIS pathway and impairs the cellular cytoprotective mechanisms.

## RESULTS

### Exposure to mild environmental stressors increases resistance to subsequent stronger challenges

In their natural habitats, animals have to cope with different types of life-threatening challenges (Fig. 1A). Prolonged exposure to environmental stressors can lead to death. For example, *C. elegans* dies within hours when exposed to strong oxidative stress or high temperature (27, 28) (Fig. 1B-C, black line). To study the interaction between different stress responses we analyzed whether a mild stress can affect *C. elegans* resistance to subsequent stronger stressors. We induced mild oxidative stress in *C. elegans* by exposing animals (L4 stage) to low non-lethal Fe^2+^ concentrations (1 h, 1mM). Fe^2+^ is a classic inductor of reactive oxygen species (ROS) in multiple biological systems, including *C. elegans* (29). After one hour of recovery, animals were subsequently exposed to a stronger oxidative stress (3 mM Fe^2+^). To quantify possible hormetic effects we calculated a survival index (SI = fraction of surviving pre-treated animals - fraction of surviving naïve animals). This value is positive when pre-exposure increases stress resistance compared to naïve animals. Animals pre-exposed to mild oxidative stress were more resistant to subsequent strong oxidative stress than naive animals (SI = 0.24 ± 0.09, Fig. 1B). Next, we analyzed if a short non-lethal exposure to high temperature (20 min, 35°C) enhanced the resistance to a subsequent prolonged exposure to the same high temperature. Similar to our observations with oxidative stress, short pre-exposure to heat increased animal survival at high temperatures (SI = 0.19 ± 0.04, Fig. 1C). Our results indicate that pre-exposure of *C. elegans* to a mild stressor enhances the animal’s resistance to subsequent exposure to the same stressor, which confirms the existence of hormesis in *C. elegans (11).*

**Fig. 1.**
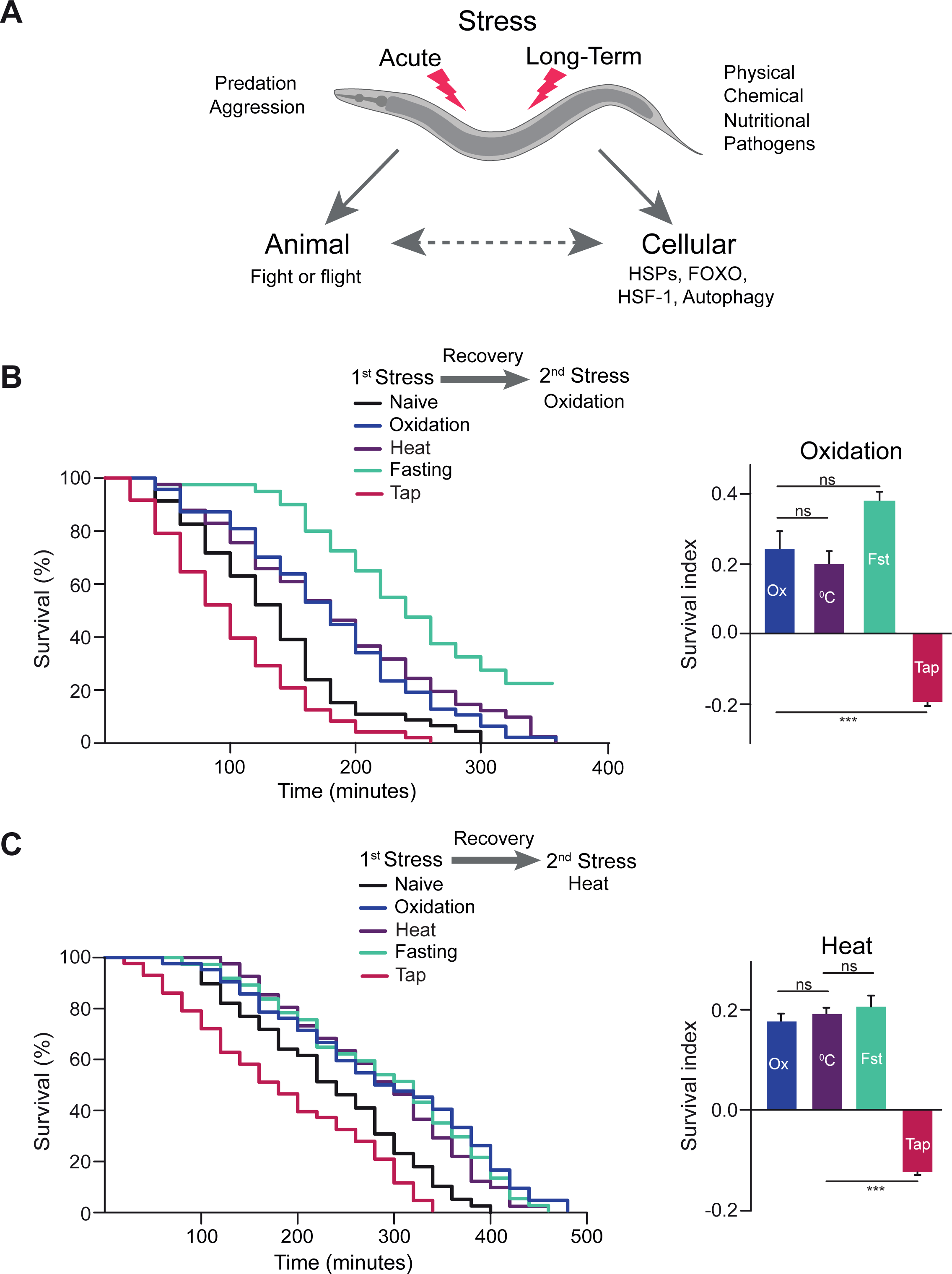
The escape response impairs the resistance to subsequent environmental stressors. **(A)** Different types of stress induce behavioral and cellular responses. (B) Scheme for the sequential stress experiments. Pre-stressors: 1 h 1 mM Fe^2+^ (Oxidation), 0.5 h 35°C (Heat), 8 h food deprivation (Fasting), or vibrational stimulus every 5 min for 2.5 h (Tap). After 1 h of recovery, survival to strong oxidative stress was evaluated. Representative Kaplan-Meier survival curves of pre-stressed nematodes exposed to subsequent strong oxidative stress (3 mM Fe^2+^). Right: Survival index (SI) at 2 h of exposure to strong oxidative stress. Positive values mean protection while negative values arise for impairment of oxidative stress resistance compared to non-treated animals. (C) Identical to B, but. heat (4 h 35°C) being the second stressor. For B and C: curves represent 3 independent replicates. Bars represent mean ± SEM, n=3-4, 35-50 worms/condition. \**p*<0.05, \*\*\**p*<0.001, One-way ANOVA followed by Holm-Sidak’s test for multiple comparisons against pre-oxidation. Summary of the lifespan experiments is presented in Table S1.

To determine whether pre-exposure with one stressor can induce crossresistance to other types of stressors, animals were briefly exposed to high temperature (20 min, 35°C) and analyzed for subsequent resistance to strong oxidative stress (Fig. 1B). Strikingly, pre-exposure to heat resulted in an increased resistance to strong oxidative stress (Fig. 1B, SI = 0.20 ± 0.06). This enhanced survival to oxidative stress is similar to that achieved after iron pre-exposure. In the reciprocal experiment animals were pre-exposed to low oxidative stress, followed by exposure to high temperature. Pre-exposure to low oxidative stress also increased thermotolerance to a similar extent to that of mild-heat pretreatment (Fig. 1C, SI = 0.18 ± 0.03). This indicates that mild heat and oxidation provide cross-resistance to subsequent stronger forms of these stressors.

Does this hormetic cross-resistance extend to other forms of stress? Another major stressor limiting animal survival, growth, and reproduction is food deprivation. To analyze whether nutritional stress can affect tolerance to either oxidative or thermal stress, animals were food-deprived for eight hours and, after recovery, exposed to either oxidative or thermal stress. Food-deprived animals were more resistant to oxidative (SI = 0.37 ± 0.05) and thermal stress (SI = 0.20 ± 0.06) compared to untreated worms (Fig. 1B and C). Therefore, our results indicate that mild chemical-, physical- and nutritional-stressors can confer cross-protection to subsequent challenges. This suggests that hormesis relies on common protective mechanisms that allow animals to better resist subsequent challenges independently of the nature of the environmental stressor.

### The *C. elegans* escape response impairs the resistance to subsequent stressors

In addition to long-term stressors, *C. elegans* also encounters acute stressors in its environment (Fig. 1A). For instance, *C. elegans* engages in a flight response to escape from predacious fungi that ensnare nematodes with hyphal nooses (26). This escape response, in which the animals reverse and turn away from the stimulus, is prompted by the activation of mechanosensory neurons (30, 31). To analyze if the escape response affects tolerance to subsequent environmental stressors, we repetitively triggered the escape response by applying a vibrational stimulus to the side of the plate (“tap”) every 5 minutes for 2.5 hours. This spaced protocol elicited an escape response after every mechanical tap (32, 33), with no obvious adverse consequences on behavior or survival (see also Fig. 3B). After a 1 hour recovery period, animals were subjected to either oxidative or thermal stress (Fig. 1B and 1C). In sharp contrast to the protective effects observed with mild environmental stressors, the repetitive induction of the escape response markedly reduced resistance to subsequent oxidative (Fig. 1B, SI = -0.19 ± 0.03) or thermal stress (Fig. 1C, SI = -0.12 ± 0.01). These results suggest that the escape response impairs resistance to subsequent long-term environmental stressors.

### Tyramine deficient animals are resistant to environmental stress

Tyramine plays a crucial role in the coordination of independent motor programs of the C. *elegans* escape response (25, 26). Since the escape response impaired resistance to subsequent environmental stressors we analyzed the role of tyramine in the animal’s environmental stress response. We found that tyramine-deficient *tdc-1* mutants are more resistant to both Fe^2+^ and H2O2-induced oxidative stress than wild-type animals (Fig. 2A and Fig. S1B-C). Furthermore, *tdc-1* mutants have increased thermotolerance compared to wild-type animals (Fig. 2A and Fig. S1D). *tdc-1* mutants have no obvious defects in pharyngeal pumping (Fig. S1E), indicating that dietary restriction is not the cause of the enhanced stress resistance. Tyramine is also a precursor for octopamine biosynthesis (22) (Fig. S1A). *tbh-1* mutants, which still can produce tyramine, but fail to synthesize octopamine, displayed normal sensitivity to oxidative stressors (Fig. S1B, C). Octopamine-deficient mutants were slightly more heat resistant than wild-type animals, albeit not at the level of tyramine/octopamine deficient *tdc-1* mutants (Fig. S1D). Moreover, rescue of *tdc-1* expression in only the octopaminergic neurons *(tdc-1; Ptbh-1::TDC* -1) failed to reduce thermoresistance of *tdc-1* mutants (Fig. S1D). These results indicate that the lack of tyramine underlies the oxidative and thermal resistant phenotype of *tdc-1* mutants.

**Fig. 2.**
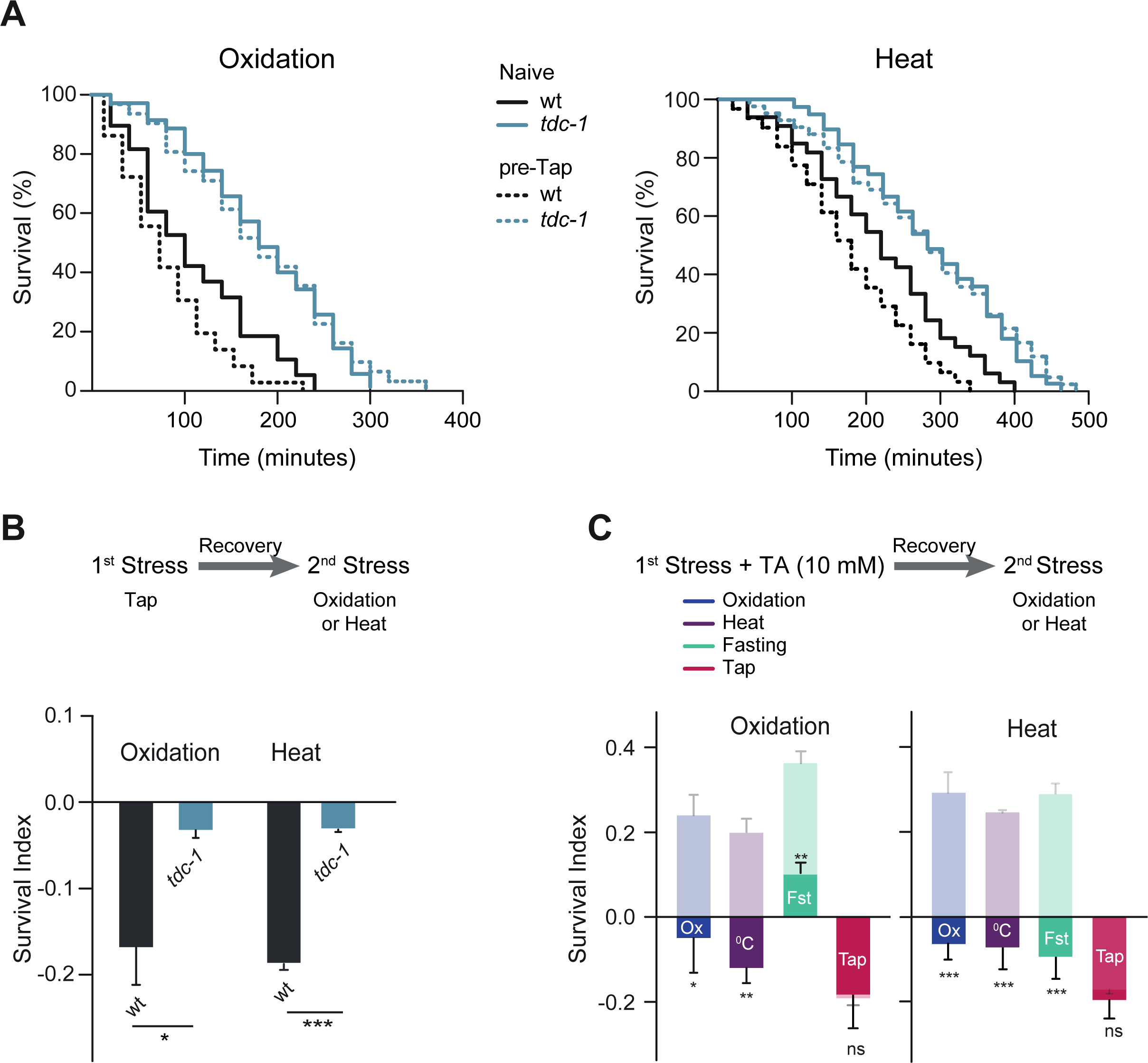
The negative effect of the escape response on stress resistance is mediated by tyramine. (A) Representative Kaplan-Meier survival curves of naïve and pre-stressed (tap) animals exposed to either oxidation (3 mM Fe^2+^, left) or heat (35°C, right)̤ Summary of the lifespan experiments are presented in Table S1. **(B)** SI of naïve and pre-stressed, wild-type and *tdc-1* mutant for oxidation and heat. Bars represent mean ± SEM, n=3, 30-45 worms/condition. * *p*< 0.05, \*\*\**p*<0.001, Student’s t-test. **(C)** Exogenous Tyramine inhibits the hormetic effects of weak environmental stressors. Pre-stressors are as described in Fig. 1 with the addition of exogenous tyramine (10 mM) during exposure to the mild stressor. SI after pre-exposure to mild stressor followed by exposure to strong oxidation (2 h, 3 mM Fe^2+^, left) or heat (4 h, 35°C, right). Bars represent mean ± SEM, n=4-5, 40-80 worms/condition) in the absence (shaded) and presence of tyramine (dark) during the pre-stress treatment, respectively.* *p*< 0.05, \*\**p*< 0.01, \*\*\**p*<0.001, Student’s t-test.

### Tyramine inhibits hormesis

Since the escape response impairs resistance to subsequent long-term environmental stressors and because tyramine-deficient animals are more resistant to stressors, we asked whether the release of tyramine during the escape response underlies the impaired resistance to subsequent stressors. To address this question, we analyzed if repeated induction of the escape response affected the subsequent oxidative and thermal stress resistance of *tdc-1* mutants. In contrast to wild-type, pre-tapped *tdc-1* animals, did not exhibit impairments in their resistance to either strong oxidative (SI = -0.03 ± 0.02) or thermal stress (SI = -0.03 ± 0.01) (Fig. 2A and B). This suggests that, tyramine release during the escape response impairs the animal’s capacity to respond to subsequent environmental stressors.

To further test our hypothesis that tyramine inhibits hormesis, we repeated the hormesis experiments in the presence of exogenous tyramine during exposure to the first minor stressor (Fig. 2C). Exogenous tyramine reversed, or strongly reduced the hormetic effects of animals exposed to mild oxidative, thermal, or nutritional pre-stressors (Fig. 2C). Pre-incubation with exogenous tyramine in the absence of environmental pre-stressors did not produce significant differences in subsequent thermal and oxidative stress resistance (Fig. S2). While tyramine signaling increases the animal’s chances to survive acute predatory challenges (26), our experiments indicate that tyramine inhibits hormesis.

### Activation of tyraminergic neurons impairs resistance to environmental stress

Acute and long-term stressors may pose conflicting challenges for the organism. Acute stressors, such as predation, trigger a rapid, energy demanding response, while prolonged environmental stressors, like starvation, require a more sustained strategy that conserves energy. What are the consequences of simultaneous activation of these opposing demands? To answer this question, we analyzed whether an animal’s response to environmental stress is affected while subjected to an acute stressor. Repeated activation of the escape response by mechanical tap, during exposure to oxidative or heat stress reduced survival of wild-type animals compared to animals that were subjected to oxidative or heat stress alone (Fig. 3, Fig. S3A). While the repeated mechanical tap by itself has no obvious consequences on viability (Fig. 3B), we could not exclude that this vibrational stimulus caused physical damage that impairs the animal’s defense mechanisms. To avoid mechanical stress on the animal, we triggered the escape response by optogenetic activation of mechanosensory neurons in transgenic animals expressing the red-shifted channelrhodopsin, ChRimson, under the *Pmec-4* promoter. Transgenic Pmec-4::ChRimson animals that were grown on plates containing the ChRimson co-factor all*-trans* retinal (ATR) showed robust behavioral responses to red light, indistinguishable from the escape response triggered by mechanical plate tap (Fig. 3D). Optogenetic activation of the escape response in Pmec-4::ChRimson animals showed reduced survival to oxidative stress and heat compared to animals raised in the absence of ATR (Fig. 3D; Fig. S3B). The mere addition of ATR, without optogenetic activation, caused no effect on the survival of animals exposed to either oxidative or thermal stress (Fig. S3C). This indicates that the activation of the escape response itself, instead of tap-induced mechanical impact, impairs resistance to environmental stressors.

**Fig. 3.**
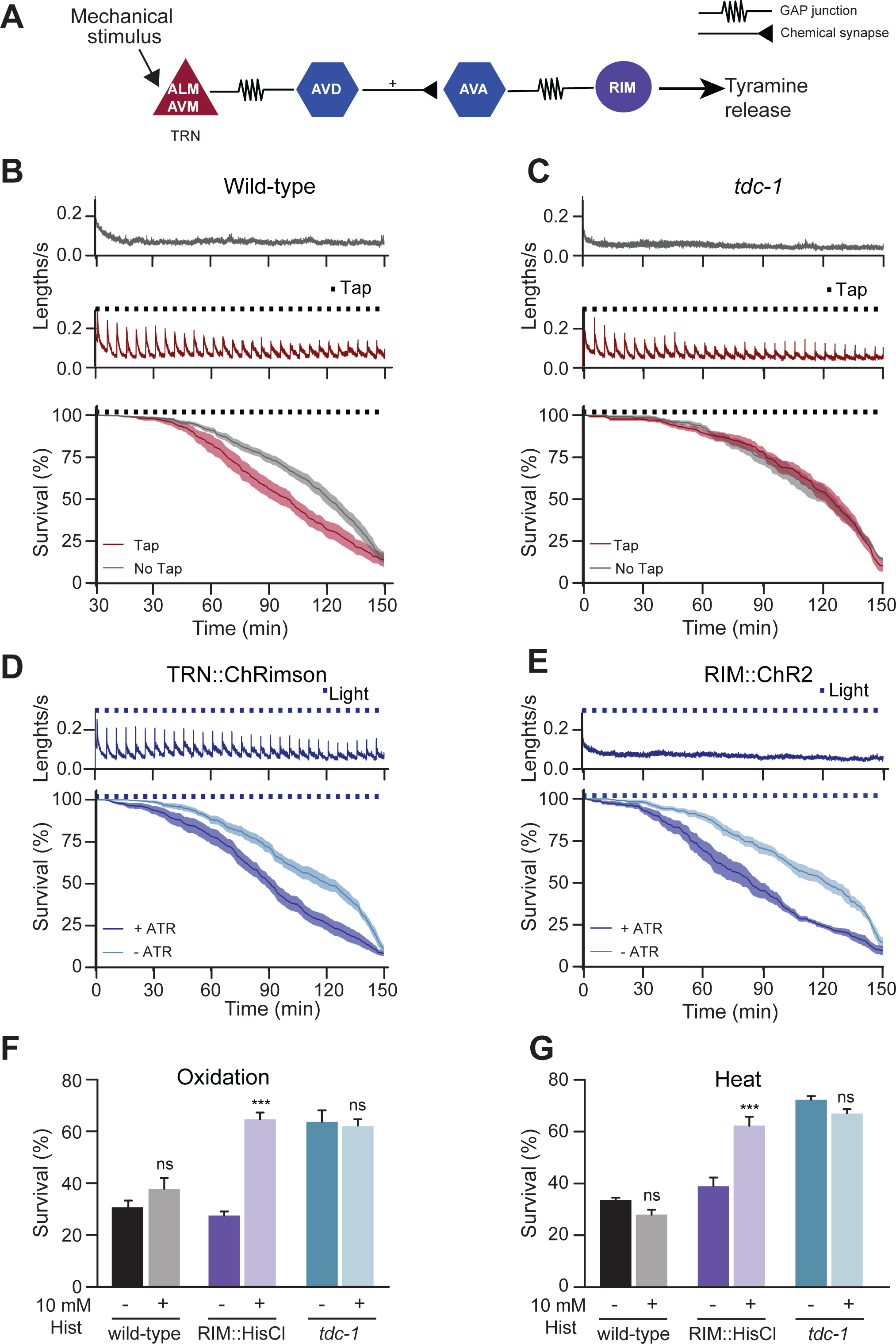
Optogenetic activation of RIM neuron is sufficient for impairing the response to environmental stressors. (A) Neural circuit for *C. elegans* escape response triggered by mechanical stimulation. **(B-C)** Velocity traces and survival curves during strong oxidative stress (3 mM Fe^2+^) for wild-type (B), or *tdc-1* mutants (C), in the absence or presence of a mechanical stimulus (tap). Velocity remains constant over the 2.5 h duration of recording in the absence of a stimulus (top), but increases rapidly in response to tap (middle). When animals are placed on Fe^2+^ plates, tap delivery reduces stress resistance in the wild-type but not in *tdc-1* mutants (bottom). Black squares indicate timing of tap delivery. Red line: tap, grey line: no tap. **(D)** Optogenetic activation of the mechanosensory neurons results in velocity increases when animals are exposed to a 5 seconds light stimulus (top) and a reduction of oxidative stress resistance (bottom). Blue line: survival curve of animals raised with all-trans retinal (ATR); grey line: animals raised without ATR. Blue squares indicate timing of light delivery. Strain used: QW1649 *zfIs144[Pmec-4::Chrimson::wCherry, pL15EK].* **(E)** Optogenetic activation of the RIM neuron does not result in velocity changes (top), but is sufficient to reduce oxidative stress resistance in animals raised with ATR (bottom). Blue squares indicate timing of light delivery. Strain used: QW1602 *lin-15(n765ts); zfEx758[Pcex-* 1::NLSwCherry::SL2::GCaMP6]. **(F-G)** Stress survival of animals expressing the histamine-gated chloride channel HisCl in the RIM neuron (RIM::HisCl). Animals were exposed to 10 mM histamine (Hist) prior and during oxidation (1 h, 15 mM Fe^2+^, F) or heat (4 h, 35°C, G). Specific silencing of the RIM neuron leads to increased animal resistance to environmental stress. Bars represent mean ± SEM, n=5, 80-100 worms/condition. \*\*\**p* < 0.001, Student’s t-test.

Previous studies proposed that stimulation of the ALM/AVM touch receptor neurons (TRN) initiates the escape response through activation of the AVD/AVA backward pre-motor interneurons (30, 31) (Fig. 3A). The AVA neurons in turn activate the single pair of tyraminergic RIM neurons (23, 34). Tyramine signaling coordinates head movements, reversal length and ventral bending during the escape response. Is activation of the tyraminergic RIM neurons sufficient to impair stress resistance? Optogenetic activation of RIM, in transgenic animals that express channelrhodopsin (ChR2) in the tyraminergic neurons *(Pcex-1*::ChR2), was not sufficient to trigger an escape response (Fig. 3E). However, RIM activation did reduce resistance to oxidative stress in animals raised on ATR compared to those lacking ATR (Fig. 3E). Furthermore, exposure to simultaneous tapping and oxidative or heat stress did not affect survival of tyramine-deficient *tdc-1* mutants (Fig. 3C, Fig. S3A). Finally, we analyzed the stress resistance of transgenic animals in which tyramine release is inhibited through the expression of a *Drosophila* histamine-gated chloride channel in the RIM neurons *(Pcex-* 1::HisCl1) (35, 36). Similar to tyramine-deficient mutants, silencing of RIM neurons by exogenous histamine increased resistance to oxidative and heat stress compared to control animals (Fig. 3F-G). These results indicate that tyramine release from the RIM neurons impairs resistance to long-term stressors.

### Dynamic changes in tyraminergic signaling underlie the response to acute *vs* long-term stressors

While tyramine signaling is beneficial during the escape response (26), tyramine signaling negatively impacts the response to environmental stress. This suggests that the up- or down-regulation from basal levels of tyramine signaling may provide a neural signal to switch between acute and the long-term stress responses. To test this possibility, we monitored calcium levels in the tyraminergic RIM neurons as a proxy for neural activity. Calcium transients were analyzed by expressing GCaMP6 in the RIM neurons of freely-moving animals that were subjected to either acute or longterm stress. In response to mechanical plate tap, there was an immediate rise in RIM calcium levels (Fig. 4A). This increase in RIM activity upon an acute stress stimulus is consistent with neuronal models of the escape response (22, 23, 34). To examine the effect of environmental stress on RIM neurons, animals were loaded into a microfluidic device and exposed to either oxidative stress or food deprivation after an initial 20-40 min acclimation period. Upon exposure to 6 mM Fe^2+^, there was a gradual but sustained reduction of calcium levels in RIM neurons, reaching maximum reduction after roughly 20 minutes of oxidative stress (Fig. 4B). Similarly, removal of food also resulted in a gradual decrease in calcium levels in RIM neurons (Fig. 4C). Conversely, when animals that were food-deprived overnight were re-exposed to food, there was a sustained increase in RIM calcium levels. This increase was initiated within 10 minutes after food addition indicating that RIM activity quickly recovers and is likely not due to GCaMP expression changes (Fig. S4). Furthermore, fluorescence levels of a RIM-expressed fluorescence marker that is insensitive to calcium (mCherry) were not decreased upon oxidation or starvation (Fig. 4B and C). This indicates that exposure to environmental stress results in the active down regulation of RIM activity and tyramine signaling.

**Fig. 4.**
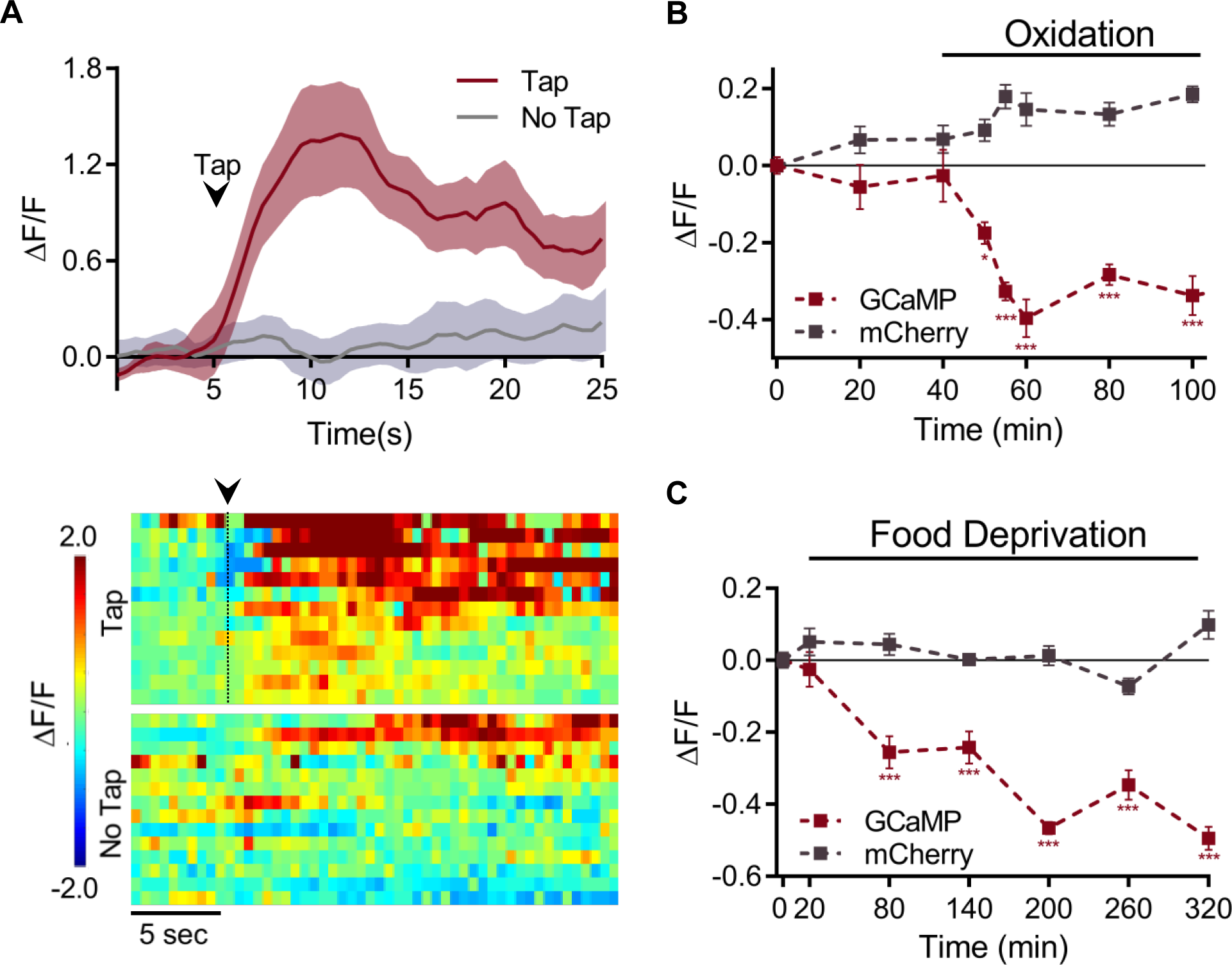
The tyraminergic RIM neuron exhibits inversed activation patterns on response to acute and gradual stressors. **(A)** (Top) Acute mechanical stress induces a rapid increase in calcium levels in RIM neurons. Traces on top represent the average RIM GCaMP6 Ca^2+^ activity (ΔF/F) 5 seconds before and 20 seconds after an mechanical stimulus to the plate in freely-moving animals (Red trace, ~1 sec tap stimulus at black arrow, 13 animals) or in control animals (Grey trace, 14 animals). Shaded areas around the traces represent SEM for the averaged traces. Bottom: Each row from the heat maps represents the GCaMP6 fluorescence from individual animals that make up the average traces. Traces were sorted by showing the highest change to the lowest change in calcium levels. **(B)** Oxidative stress (6 mM Fe^2+^) reduces calcium levels in RIM neurons. Overall RIM calcium levels (ΔF/F) are significantly decreased after exposure to oxidation (red trace, 36 animals/condition, n=6). Oxidation does not affect mCherry fluorescence intensity in RIM neurons (grey trace, 15 animals/condition, n=5). *p*<0.0001, one-way ANOVA, \**p*<0.05, \*\*\**p*<0.001 compared to initial time point, Dunnett’s multiple comparison. (C) Food-deprivation reduces Ca^2+^ levels in RIM neurons. Overall RIM Ca^2+^ levels (ΔF/F) are significantly decreased after food deprivation (red trace, 30 animals, n=5). Food-deprivation does not affect mCherry fluorescence intensity in RIM neurons (grey trace, 6 animals/condition, n=2). *p* <0.0001, two-way ANOVA, *** *p* < 0.001, compared to initial time point, Dunnett’s multiple comparison.

### Activation of adrenergic-like TYRA-3 receptor inhibits the long-term stress response

What are the targets of tyramine? *C. elegans* has four different receptors that are activated by tyramine: three adrenergic-like GPCRs (SER-2, TYRA-2 and TYRA-3) (37) (38) (39) and a tyramine-gated chloride channel (LGC-55) (23, 40). We analyzed mutants for each of these receptors to determine which tyramine receptor(s) underlies the effect of tyramine on stress resistance. Like tyramine-deficient animals, *tyra-3* mutants were resistant to oxidative stress, heat and starvation (Fig. 5A; Fig. S6). Other receptor mutants did not display obvious increases in resistance. Moreover, exogenous tyramine, which impairs the stress resistance of wild-type and *tdc-1* animals (Fig. 5B), did not reduce the stress resistance of *tyra-3* mutants (Fig. 5B). This shows that tyraminergic activation of TYRA-3 is required to modulation the stress response.

**Fig. 5.**
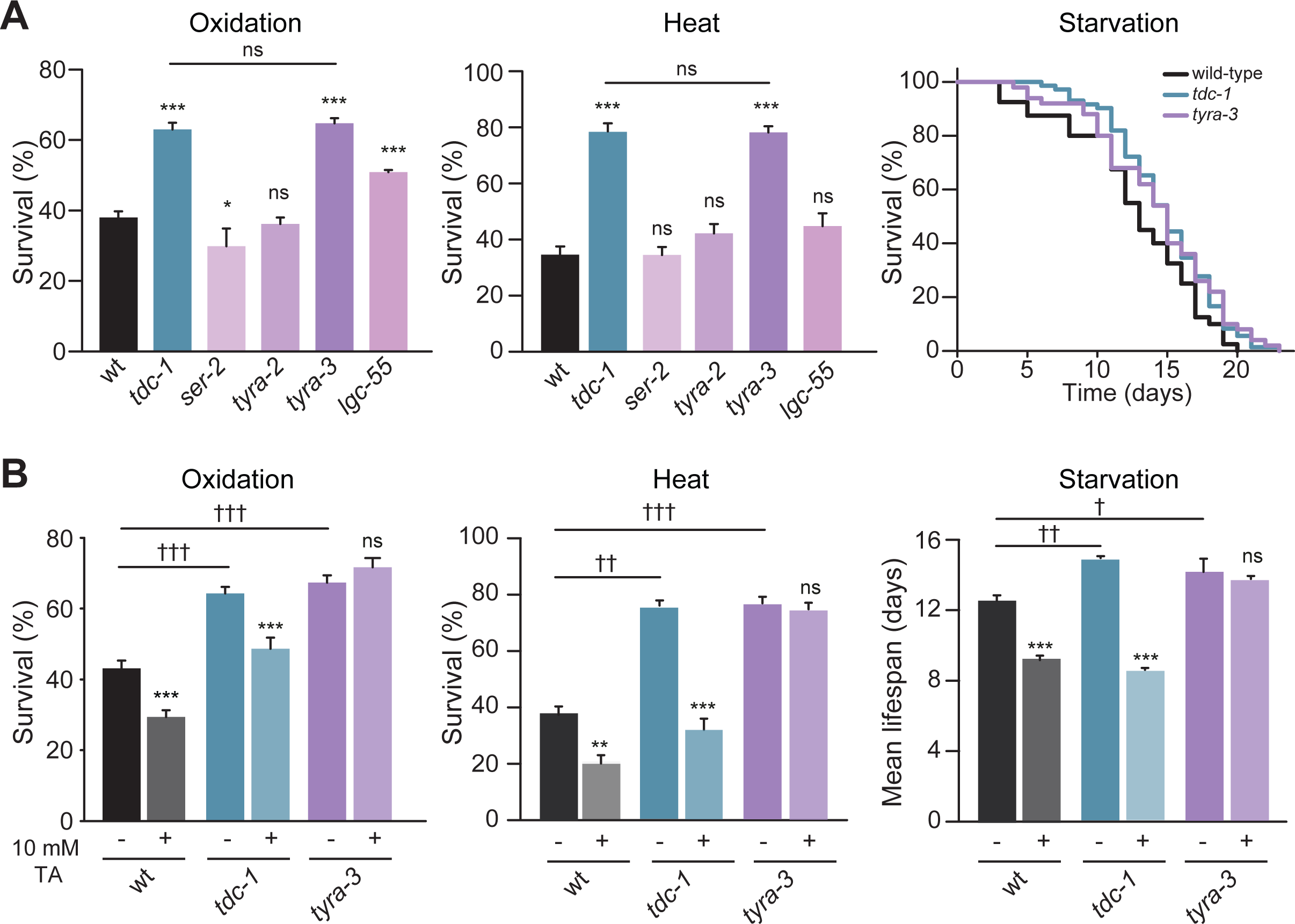
Tyraminergic coordination of the stress response is performed through the activation of the GPCR TYRA-3. (A) Resistance of wild-type and tyramine receptor null-mutants exposed to oxidation (1 h, 15 mM Fe^2+^) or heat (4 h, 35 °C). Only *tyra-3* mutant animals are as resistant as *tdc-1* mutant animals. Bars represent mean ± SEM, n=3-6, 80-100 animals/condition. \**p*<0.05, \*\*\**p*<0.001, One-way ANOVA followed by Holm-Sidak’s test for multiple comparisons. (Right) Representative Kaplan-Meier survival curves of wild-type, *tdc-1* and *tyra-3* mutants exposed to starvation. Curves represent 3 independent replicates. Summary of the lifespan experiments are presented in Table S1. **(B)** Resistance of wild-type, *tdc-1* and *tyra-3* mutant animals exposed to oxidation, heat or starvation in the absence and presence of exogenous tyramine (10 mM). Detrimental effects of exogenous tyramine on stress resistance are abolished in *tyra-3* mutant animals. Bars represent mean ± SEM, n=3-4, 60-80 worms/condition. † *p*<0.05, †† *p*<0.01, ††† *p*<0.001 compared to wild-type and \*\**p*<0.01 and \*\*\**p*<0.001 compared to absence of tyramine within each strain, two-way ANOVA, followed by Holm-Sidak’s test for multiple comparisons.

Where does TYRA-3 act? A *Ptyra-3::GFP* reporter, driven by a 3.6 kb promoter is expressed in a subset of neurons in the nervous system and the intestine (Fig. 6A). We identified a 1.8 kb *tyra-3* promoter fragment, which drives reporter expression in the neuronal subset, but not the intestine *(Ptyra-3*_*short*_*::mCherry;* Fig. S5). Expression of *tyra-3* in neurons *(Ptyra-3*_*short*_*::TYRA-3)* failed to rescue the stress resistant phenotype in *tyra-3* mutants. However, expression of *tyra-3* in the intestine (Pelt-2::TYRA-3) was sufficient to restore the stress sensitivity of *tyra-3* mutants to wild-type levels (Fig. 6B-D, Fig. S6). Intestinal expression of *tyra-3* also restored the negative impact of exogenous tyramine on heat, oxidative stress and starvation survival assays (Fig. S6). Furthermore, like tyramine deficient mutants, repeated induction of the escape response in *tyra-3* mutants did not increase sensitivity to subsequent heat or oxidative stress (Fig. 6D). The tap-induced increase in sensitivity to environmental stressors was restored when *tyra-3* is expressed in the intestine (Fig. 6D). Since the RIM neuron has no direct synaptic outputs onto the intestine, these experiments show that tyramine acts as a neurohormone to inhibit the long-term stress response through the activation of TYRA-3 in the intestine.

**Fig. 6.**
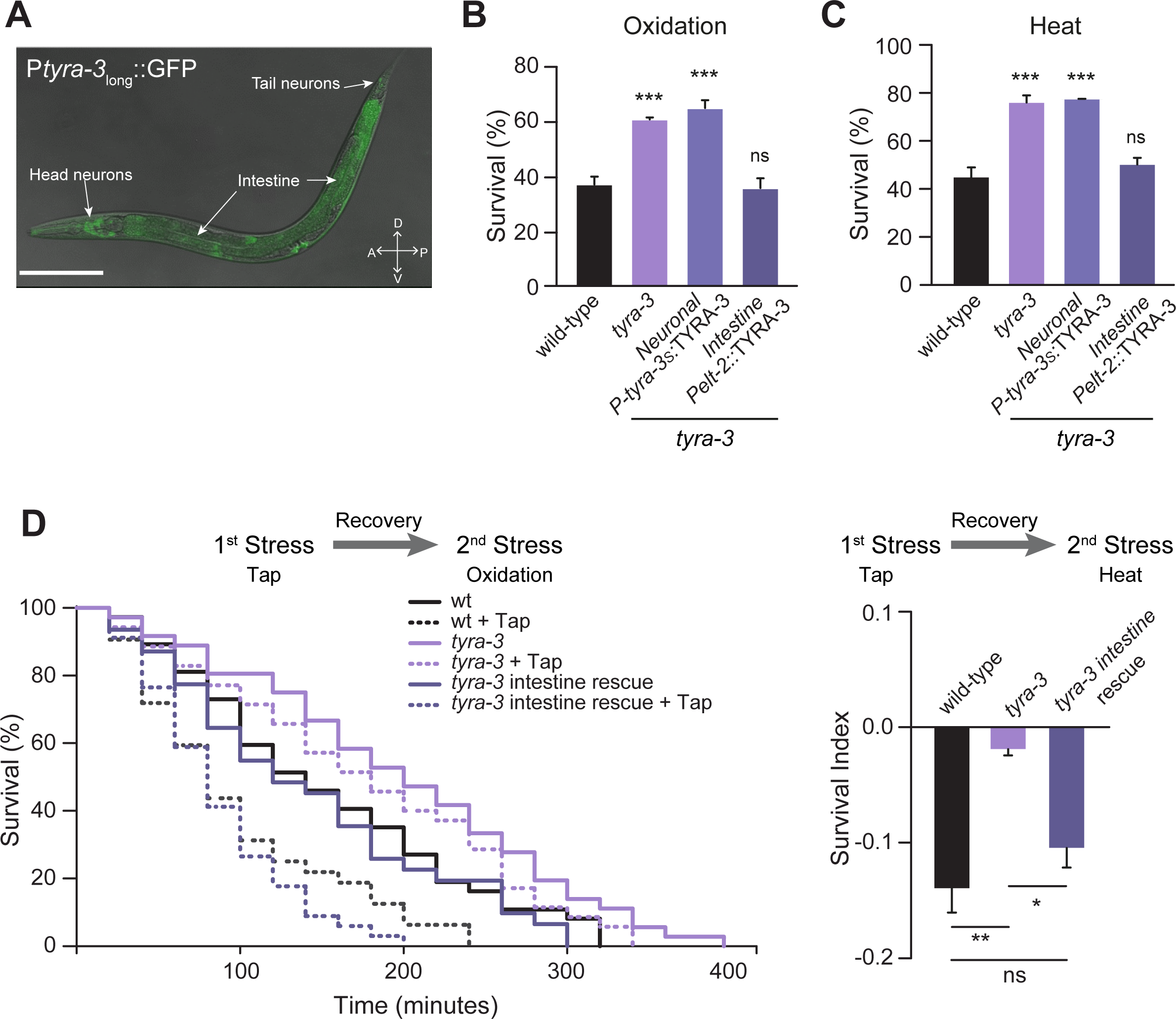
TYRA-3 intestinal expression is essential for the tyraminergic modulation of the stress response. (A) Representative confocal fluorescence image of transgenic animals expressing GFP driven by a 3.6 kb promoter of *tyra-3a (Ptyra-3long::GFP).* GFP expression is observed in both nervous system and intestine. Scale bar: 150 μm. **(B-C)** Survival percentages of *tyra-3* null mutant animals expressing either the neuronal *(Ptyra-3s::TYRA-3)* or the intestinal *(Pelt-2::* TYRA-3) rescue constructs exposed to oxidative stress (1 h, 15 mM Fe^2+^) (B) or heat (4 h, 35°C) (C). Bars represent mean ± SEM, n=3-6, 80-100 animals/condition). \*\*\**p*<0.001, One-way ANOVA followed by Holm-Sidak’s test for multiple comparisons. (**D**) Left: Representative Kaplan-Meier survival curves of naïve (solid line) or pre-exposed to tapping (dashed line) exposed to strong oxidative stress (3 mM Fe^2+^). Curves represent 3 independent replicates. Summary of the lifespan experiments are presented in Table S1. Right: SI to heat exposure (4 h, 35°C) of animals pre-exposed to tapping. Intestinal expression of *tyra-3* restores the detrimental effect of tapping on the stress response. Bars represent mean ± SEM, n=3, 30-40 animals/condition. \*\**p*<0.01, \**p*<0.0.5, Oneway ANOVA followed by Holm-Sidak’s test for multiple comparisons.

### Tyraminergic inhibition of systemic stress response depends on the DAF-2/IIS pathway

The insulin/IGF-1 signaling (IIS) pathway regulates growth, reproduction, metabolic homeostasis, lifespan and stress resistance from nematodes to humans (41, 42). In *C. elegans*, loss-of-function mutations in the insulin/IGF-1 receptor ortholog, DAF-2, increase stress resistance and longevity (43, 44). To test whether tyramine acts through the DAF-2/IIS pathway, we analyzed the effect of tyramine signaling on the stress resistance of *daf-2* mutants. Stress resistance of *tdc-1; daf-2* and *tyra-3; daf-2* double mutants was similar to that of *daf-2* single mutants (Fig. 7A). Furthermore, exogenous tyramine did not impair the stress resistance of *daf-2* mutants (Fig. S7). This suggests that tyramine inhibition of the stress response is dependent on DAF-2/IIS pathway.

**Fig. 7.**
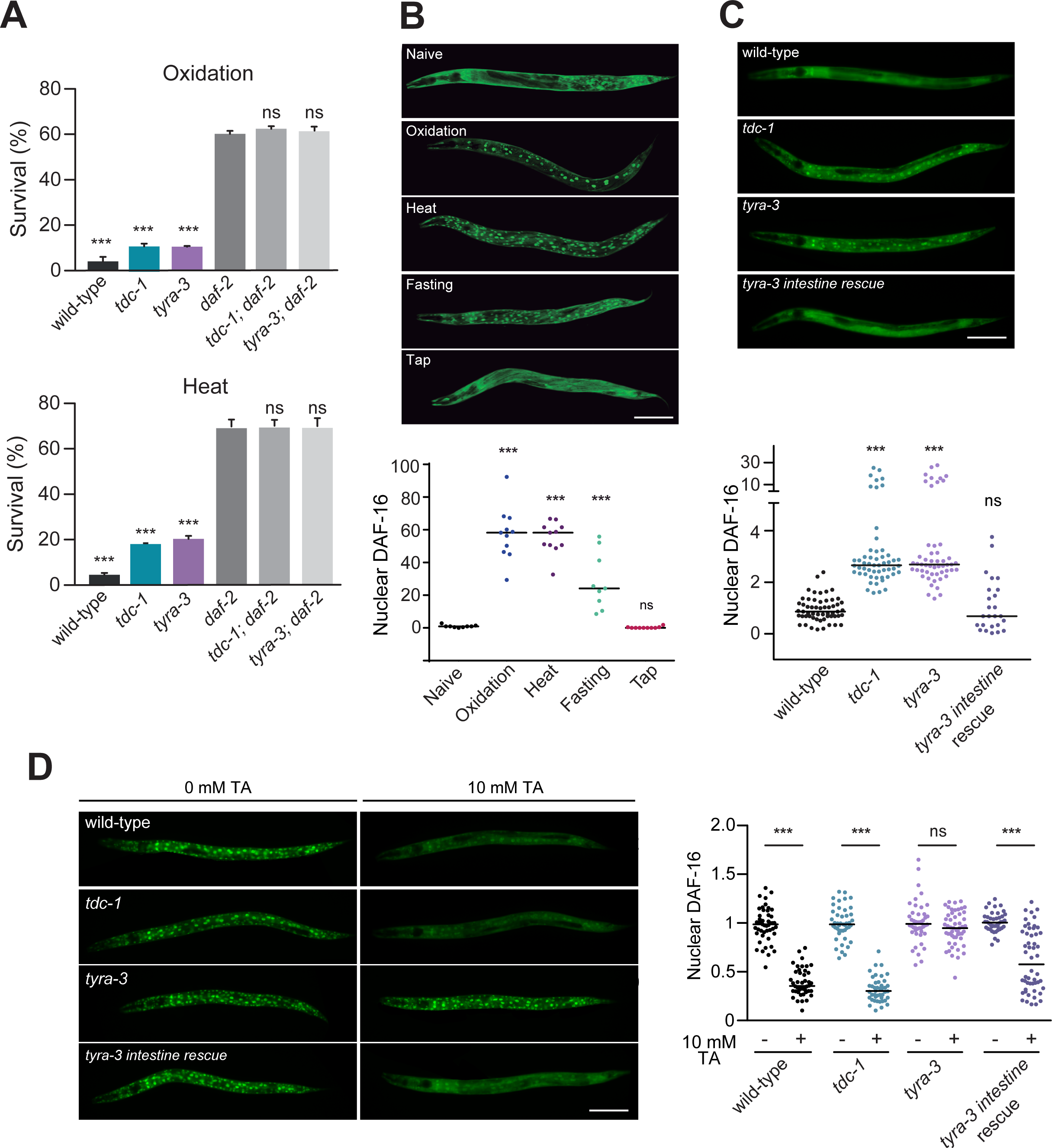
Tyramine signaling inhibits stress-dependent nuclear translocation of DAF-16. **(A)** Survival percentage of animals exposed to oxidative stress (3 h, 15 mM Fe^2+^, top) or heat (7 h, 35°C, bottom). *tdc-1; daf-2* and *tyra-3; daf-2* double mutants are as resistant as *daf-2* single mutant worms. Bars represent mean ± SEM, n=3-4, 80 animals/condition. \*\*\**p*<0.001 compared to *daf-2*, one-way ANOVA followed by Holm-Sidak’s test for multiple comparisons. (**B**) Top: Representative fluorescence images of worms expressing the translational reporter *Pdaf-16::DAF-16a/b::GFP* after exposure to mild stressors. Scale bar: 150 μm. Bottom: Scatter dot plot (line at the median) with the number of nuclear DAF-16 per worm (normalized to naïve animals). Nuclear localization of DAF-16 is significantly higher in animals exposed to long-term stressors compared to animals exposed to an acute stressor (tap). 10-15 worms/condition. \*\*\**p*<0.001, compared to naïve, one-way ANOVA followed by Holm-Sidak’s test for multiple comparisons. (C) Top: Representative fluorescence images after mild heat exposure (10 min, 35°C). Scale bar: 150 μm. Bottom: Scatter dot plot (line at the median) with the number of nuclear DAF-16 per worm (normalized to wild-type animals). Nuclear localization of DAF-16 is significantly higher in *tdc-1* and *tyra-3* mutants. 30-60 animals/condition. \*\*\**p*<0.001, compared to wild-type, One-way ANOVA followed by Holm-Sidak’s test for multiple comparisons. (**D**) Left: Representative fluorescence images (20x magnification) after stronger heat exposure (30 min, 35°C) in the absence or presence of exogenous tyramine (10 mM). Animals are shown with the anterior region to the left. Scale bar: 150 μm. Right: Corresponding quantification (normalized to untreated-animals). Exogenous tyramine decreases DAF-16 nuclear localization in the wild-type, *tdc-1* mutants and animals expressing *tyra-3* solely in the intestine. \*\*\**p*<0.001, compared to untreated-worms, One-way ANOVA followed by Holm-Sidak’s test for multiple comparisons.

**Fig. 8.**
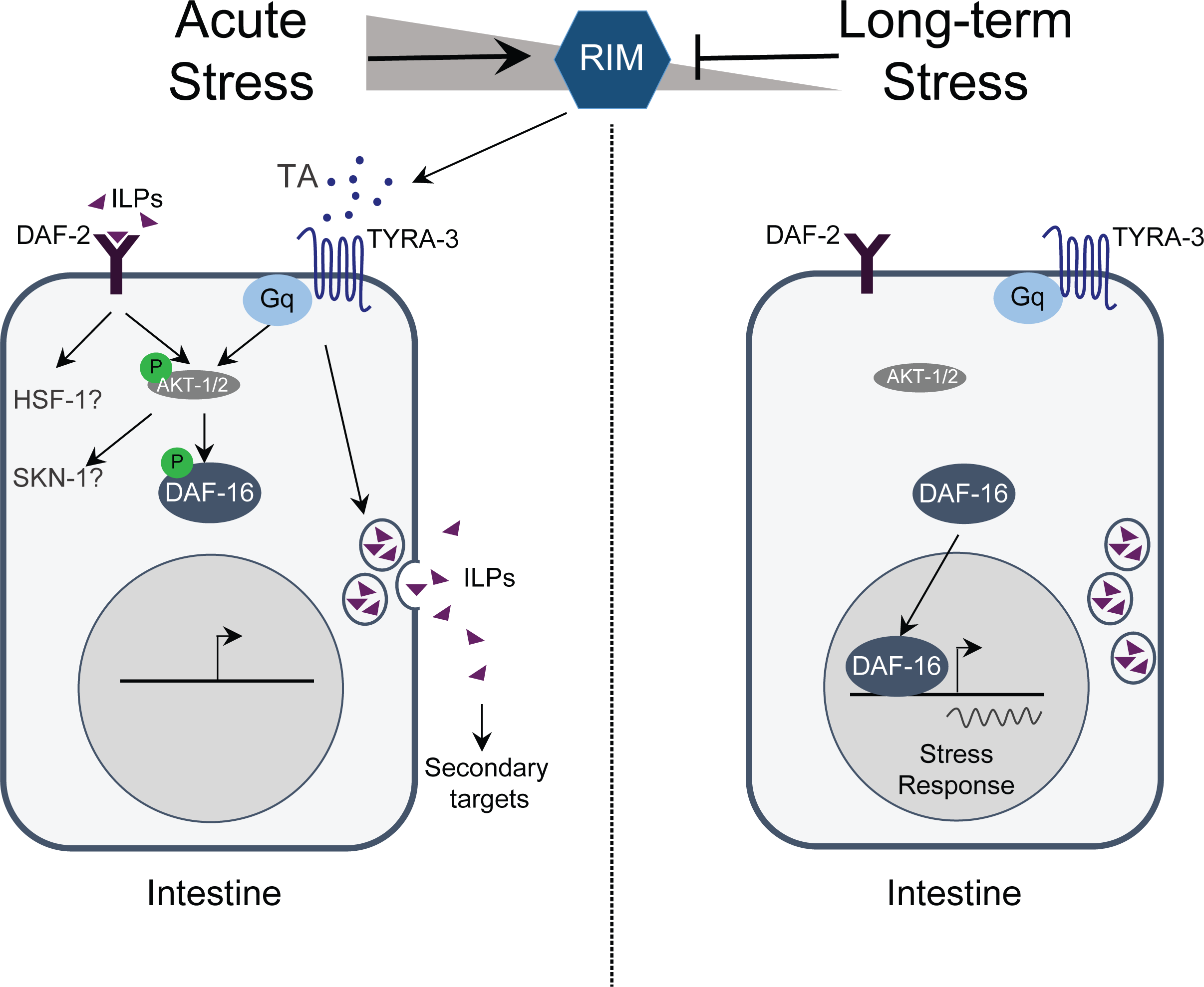
Model: Tyraminergic modulation of the DAF-2/IIs pathway. Left: Exposure to acute stressors triggers the escape response and the release of tyramine from the RIM neuron. Tyramine activates the adrenergic-like Gq-coupled receptor TYRA-3 in the intestine. Gq-protein coupled receptor signaling activates the IIS pathway. As a result, DAF-16 (and, probably, other DAF-2/IIS dependent factors such as HSF-1 and/or SKN-1) translocation to the nucleus is prevented. In parallel, TYRA-3 activation may also stimulate the release of ILPs that can activate the DAF-2 receptor in the intestine and other tissues. Right: Upon exposure to long-term stressors tyramine release from the RIM is inhibited. TYRA-3 becomes inactive leading thus no longer stimulating the DAF-2/IIS pathway. This allows for nuclear translocation of DAF-16/FOXO resulting in the transcription of cellular stress response genes.

The transcription factor DAF-16/FOXO mediates a large portion of the physiological processes that are controlled by the DAF-2/IIS pathway. Environmental stressors reduce the activity of the DAF-2/IGFR, leading to the dephosphorylation and activation of DAF-16/FOXO. Upon activation, DAF-16/FOXO translocates to the nucleus where it induces the expression of stress response genes (45, 46). In *C. elegans*, the inhibition of DAF-2/IIS signaling and nuclear translocation of DAF-16/FOXO has been described for a variety of environmental stressors (6, 47). The striking cross-resistance between heat, oxidative stress and starvation we observed in hormesis experiments (Fig. 1) could be explained by the protective activation of a common defense mechanism. Indeed, DAF-16/FOXO localized to the nucleus in response to a short exposure to heat, oxidative or nutritional stress (Fig. 7B). However, DAF-16/FOXO largely localized to the cytoplasm in response to repeated tap stimulus. To determine how the induction of the escape response and tyramine release affect the DAF-2/IIS pathway, we compared the localization of the translational reporter DAF-16::GFP in wild-type, *tdc-1* and *tyra-3* mutant animals. Under normal growth conditions DAF-16 is mostly cytoplasmic and no differences in its subcellular localization were detected (Fig. S7B). However, after a short heat exposure (35°C, 10 min), *tdc-1* and *tyra-3* mutants exhibited significantly higher levels of DAF-16/FOXO nuclear accumulation when compared to wild-type background (Fig. 7C). Upon extended heat exposure, DAF-16/FOXO (35°C, 35 min) is predominantly localized to the nucleus in both the wild-type and tyramine signaling mutants. Nevertheless, exogenous tyramine inhibits DAF-16/FOXO nuclear localization in wild-type and *tdc-1* animals, but not in *tyra-3* mutant animals (Fig. 7D). Intestinal expression of *tyra-3* rescued the DAF-16/FOXO localization phenotype of *tyra-3* mutants (Fig. 7D). These observations are consistent with the notion that tyramine signaling inhibits DAF-16/FOXO phosphorylation, and the activation of stress response genes.

## DISCUSSION

In this study we show that the acute stress response reduces worm ability to cope with environmental challenges. Acute stressors trigger the release of tyramine, which stimulates the DAF-2/IIS pathway through the activation of the adrenergic-like receptor TYRA-3 in the intestine. Exposure to long-term stressors reduces tyramine signaling, allowing DAF-16/FOXO translocation to the nucleus and induction of cytoprotective mechanisms. Tyramine signaling provides a state dependent neural signal that regulates the switch between the acute and long-term stress responses.

### Hormesis

Adaptive responses to low levels of environmental challenges have been shown to increase the resistance to stronger stressors from bacteria to plants to animals (48). If the first signs of stress are indications of worse to come, the protective activation of cellular defense mechanisms may improve stress tolerance under ensuing adversity. This phenomenon, called hormesis, has also been reported for *C. elegans (11, 49).* In accordance, we show that low-dose stressors confer protection to stronger versions of the same stressor. Moreover, we find that a low-dose of one stressor also confers cross-protection to other types of environmental stressors. The occurrence of cross-resistance indicates that cytoprotective and restorative intracellular processes are shared among different environmental stressors (50, 51). Either brief food-deprivation, low-dose oxidative or heat stress, lead to the inactivation of DAF-2/IIS, and the translocation of DAF-16/FOXO to the nucleus. DAF-16 nuclear translocation activate stress response genes required for tolerance to heat, radiation, oxidative damage, anoxia as well as pathogen infection (52-54). The shared involvement of the DAF-2/IIS pathway in response to a wide variety of harmful challenges explains, at least in part, the cross-hormetic effect of environmental stressors. Cross-resistance may be a byproduct of the overlapping effects of different stressors. For example, osmotic, ionizing stress and heat can produce oxidative damage (55-57). Alternatively, crossresistance may have evolved to prepare organisms for impending stress if successive stressful environments are frequently encountered in nature such as heat and UV radiation and desiccation.

### Neural modulation of the acute and the long-term stress response

The protective effect of the activation of systemic defense mechanisms is long lasting and can even extend the lifespan of the organism (41, 42). In contrast, acute stressors require actions on a much shorter time scale. Fight-or-flight responses against a predatory attack or aggression must occur within seconds to avoid imminent harm or even death. In *C. elegans* touch can elicit a flight- or escape-response where the animal reverses and quickly turns away from the mechanical stimulus. In sharp contrast to the hormetic cross-resistance of environmental stressors, the repetitive induction of the escape response impairs animal survival to subsequent stressors. Tyramine, which is structurally and functionally related to vertebrate adrenaline, plays a crucial role in the escape response (23, 26). While the activity of the tyraminergic RIM neuron rapidly increases during the escape response, RIM activity decreases upon exposure to environmental stressors such as oxidative stress or food deprivation. These findings support computational models that predict inhibition of RIM activity upon food deprivation (58) and are consistent with decreased RIM activity in animals exposed to pathogens (36). Furthermore, optogenetic activation of the RIM reduces, whereas RIM silencing increases resistance to environmental stressors. This indicates that while tyramine release is needed for the escape response to avoid predation (26), the down-regulation of tyraminergic signaling is required to deal with long-term stressors. Tyramine may thus provide a neural switch for the animals’ response to acute *vs* long-term stressors. Although it is currently unclear how sensory perception of environmental stressors leads to the inhibition of the RIM neuron, it is worth noting that RIM receives synaptic inputs from AIB and AIY neurons, key players in the C. *elegans* starvation- and thermal-response (20, 59, 60).

### Tyramine stimulates the DAF-2 insulin pathway

We show that tyramine acts as a neurohormone that inhibits the systemic stress response through the activation of the adrenergic-like GPCR, TYRA-3, in the intestine. During the acute stress response TYRA-3 stimulates the DAF-2/IIs pathway and inhibits DAF-16/FOXO translocation to the nucleus (Fig. 8). Stimulation of DAF-2/IIs pathway is linked to increased metabolic rate (61, 62) reviewed in (63). The metabolic rate also rises during the flight response in many animals (64, 65). Therefore, the extra-synaptic activation of TYRA-3 may induce a metabolic shift to provide the fuel needed for the high-energy demand of the C. *elegans* escape response. TYRA-3, a predicted Gq-coupled GPCR (66), may promote phospholipase C (PLC) activity and stimulate PKC and inositol 1,4,5-trisphosphate (IP3)-mediated intracellular calcium release (67). Gq-coupled adrenergic receptors have been shown to stimulate the PKC-PI3K-AKT signaling pathways (68, 69). Further studies are needed to elucidate if TYRA-3 promotes DAF-16 phosphorylation in the intestine by direct stimulation of AKT. It is noteworthy that TYRA-3 activation inhibits DAF-16 translocation, not only in the intestine, but also in other tissues. This suggests that tyramine-dependent Gq activation may also stimulate the release of Insulin-Like Peptides (ILPs) from the intestine to further inhibit cytoprotective pathways in non-intestinal cells. The fact that activation of Gq-receptors is critical for insulin secretion in mammals supports this idea (70, 71).

### Acute stress and cytoprotection

While isolated cells have the intrinsic capacity to respond to stress (e.g. HSPs, SOD and/or autophagy induction), the coordination of these cytoprotective mechanisms in multi-cellular organisms depends on the neural perception of the stressors (72). In *C. elegans* for instance, the systemic heat shock response is regulated by the AFD thermosensory neuron, which stimulates the systemic activation of HSF1 in remote tissues (7, 73). Although the activation of cytoprotective pathways safeguard against the potentially greater costs of repair, replacement, and death, it drains resources from animal development and reproduction. For instance, *C. elegans* stress resistant mutants display slow developmental rates and reduced brood sizes (74, 75). An animal benefits most from limiting the activation of cytoprotective pathways when they are not necessary, while retaining inducible capacity for emergency defense. Tyramine thus provides a neural signal to inhibit the activation of cell-intrinsic defenses. While tyramine release facilitates and prepares the animal’s escape from a threatening stimulus, the down regulation of tyramine signaling is crucial to deal with environmental stressors in which a behavioral response fails to alleviate the exposure to the stressor.

In vertebrates, acute stress stimulates the release of adrenaline and noradrenalin: key inducers of the animal’s fight-or-flight response (1, 2). Frequent exposure to acute stressors, such as predators or even predator odors leads to the chronic release of these stress hormones (76, 77). Chronic activation of the fight-or-flight response weakens the animals defense systems to other stressors (78, 79). For instance, salmon previously exposed to predators had significantly higher mortality when exposed to an osmotic stressor (80) and sparrows subject to frequent nest predation exhibit lower antioxidant capacity (79). In humans, persistent experience of either real or perceived danger has been associated with adverse health, and premature death (81, 82) (17). For example, patients that suffer from post-traumatic stress disorder (PTSD) exhibit high risks of neurodegenerative, cardiovascular, respiratory and gastrointestinal diseases, immune dysfunction and premature aging (83-86). Severity of PTSD symptoms correlates with high levels of noradrenaline, suggesting that increased sympathetic arousal may be closely linked to the dysregulation of body fitness (87-89). How increased levels of stress neurohormones impair general health and defense mechanisms in humans is currently unclear. Given the striking conservation in neural control of stress responses, it will be interesting to determine whether adrenaline and noradrenaline negatively impact health and aging through the inhibition of insulin-dependent cytoprotective pathways.

## Acknowledgments

Some strains were provided by the CGC, which is funded by NIH Office of Research Infrastructure Programs (P40 OD010440). The CX16663 strain was kindly provided by C. Bargmann. We like to thank Andres Garelli, Alex Byrne and Marian Walhout for critically reading the manuscript. We also thank Dirk Albrecht for microfluidic devices and help with the calcium-imaging set-up, Christoph Weist and Manu Madhav for analysis software, Andrea Thackeray and William Joyce for technical support and Claire Benard for DNA constructs. **Funding**: This work was supported by Grants from UNS (PGI: 24/B216 DR, PGI24/ZB62 MJDR), ANPCYT, (PICT 2014 3118 DR) and CONICET (PIP11220150100182œ DR-MJDR) and grant GM084491 from the National Institutes of Health. **Author contributions**: MJDR, TV, JF, JG, GB, NA, JD and DR performed the experiments and analyzed the data. MJDR, TV, JF, JG, DR and MJA designed the experiments. DR and MJA conceived the study, and wrote the paper. **Competing interests**: Authors declare no competing interests. **Data and materials availability**: All data is available in the main text or the supplementary materials

## Supplementary Materials

Materials and Methods

Figures S1-S7

Table S1

References (*90-97*)

